# Distinct patterns of multisensory integration underly the expression of flavor preferences and aversions in rats

**DOI:** 10.64898/2026.11.03.764007

**Authors:** Isabella B. Allar, Anna Luisa Berenguer-Nascimento, Bridget N. McNally, Yiwei Tang, Nazlee Harunani, Joost X. Maier

**Author notes:** Correspondence: Joost X. Maier, Wake Forest University School of Medicine, Department of Translational Neuroscience, Medical Center Blvd, Winston-Salem, NC 27157, USA. These authors contributed equally (Isabella B. Allar, Anna Luisa Berenguer-Nascimento).

## Abstract

Flavor is an inherently multisensory experience, combining gustatory and retronasal odor signals during consumption. Previous work has shown that rats weight taste and smell components of flavor mixture solutions to express flavor preferences. However, a comprehensive quantitative understanding of taste-smell integration is lacking. Here, we evaluated the validity of the maximum likelihood estimation (MLE) framework for explaining multisensory integration of taste and smell signals toward hedonic flavor judgments. MLE is a dominant framework for explaining multisensory integration in a wide variety of contexts, and states that components of a mixture are weighted according to their relative reliability, resulting in more reliable multisensory judgments. In our experiment, rats expressed preferences for taste, odor and taste+odor solutions in a series of two-bottle preference tests featuring an unpalatable odor component (amyl acetate), and either a palatable taste component (sucrose) or an unpalatable taste component (citric acid). Taste reliability was experimentally varied by presenting taste components at two different concentrations. Reliability of animals’ preference judgements was quantified as the variance in preference across repetitions of the same conditions in the same animals. Our results confirm that preferences for mixtures featuring a palatable taste can be described as a weighted average of the component preferences where the weights depend on the relative reliability of the components—a finding that is in line with the MLE framework. However—contrary to MLE—component weighting does not increase the reliability of multisensory preference judgments themselves. In addition, preferences for taste+smell mixtures featuring an unpalatable taste follow a qualitatively different pattern: taste and smell components are combined to enhance flavor rejection judgements. These results demonstrate that the multisensory integrative principles underlying flavor preference differ from other multisensory domains and depend on the valence of the flavor components.

## Introduction

Multisensory integration—the ability to combine multiple sensory signals associated with an object or event into a unified percept—greatly enhances subjective experience, decision making and learning (Barry E. Stein, 2012). Flavor is the quintessential example of a multisensory experience. During consumption, taste molecules stimulate the lingual epithelium, giving rise to the basic taste qualities sweet, bitter, sour, salty and umami (gustation) (Breslin & Spector, 2008). At the same time, volatile molecules released inside the oral cavity travel through the nasopharynx and stimulate the olfactory epithelium to give rise to olfactory qualities (retronasal olfaction) (Bartoshuk, Sims, Colquhoun, & Snyder, 2019; Rozin, 1982). Together with sensations of mouthfeel, these sensory qualities form the core components of flavor (Small & Green, 2012). Multisensory aspects of flavor are combined by the brain to create a coherent percept and inform perceptual judgements that contribute to ingestive behaviors (Spence, 2015). Indeed, previous research has shown that humans and other animals integrate multisensory flavor signals to inform hedonic judgments about food (Amsellem & Ohla, 2016; Colbert, Triplett, & Maier, 2022; Elliott & Maier, 2020; Fondberg, Lundström, Blöchl, Olsson, & Seubert, 2018; Hua, Dong, & Maier, 2024; Maier & Elliott, 2020; Schifferstein & Verlegh, 1996; Seo et al., 2013).

Recent work has started to elucidate the principles underlying integration of multisensory flavor signals. Behavioral studies in rats demonstrate that hedonic judgments of taste-smell (Elliott & Maier, 2020; Maier & Elliott, 2020) and taste-texture (Colbert et al., 2022) mixtures can be described as a weighted average of the unisensory component judgements. Moreover, animals vary the weight each component carries toward multisensory judgements in an adaptive, context-dependent manner, such that more ambiguous sensory cues carry less weight than more reliable ones (Allar, Hua, Rowland, & Maier, 2025; Maier & Elliott, 2020). These findings are roughly consistent with the maximum likelihood estimation (MLE) framework for explaining multisensory judgements in other systems, including visual-haptic judgements of object size (Ernst & Banks, 2002), auditory-visual judgments of object location (Alais & Burr, 2004; Meijer, Veselič, Calafiore, & Noppeney, 2019) and visual-vestibular perception of heading direction (Butler, Smith, Campos, & Bülthoff, 2010; Fetsch, Pouget, DeAngelis, & Angelaki, 2011; Fetsch, Turner, DeAngelis, & Angelaki, 2009). According to this framework, multiple sensory cues contribute to judgements of a single stimulus quality in a statistically-optimal manner. Specifically, the MLE framework makes two predictions regarding perceptual judgements of multisensory stimuli: 1) Multisensory judgements are a weighted average of the component judgements, and the weights carried by the components are determined by relative reliability where reliability is measured by the variance in judgement; and 2) The resulting multisensory judgements are more reliable (i.e., less variable) than the component judgements.

A previous study from our lab supports the first prediction of the MLE framework with respect to hedonic flavor judgments. Taste and smell components of a flavor mixture are weighted according to their relative reliability to inform preference judgements (Allar et al., 2025). The second prediction remains untested in the context of flavor preference judgements. Flavor preference judgments also differ markedly from the type of perceptual judgements used in previous work. In typical psychophysical studies, subjects make judgements of individual stimuli on a trial-by-trial basis. In contrast, flavor preferences are derived from consumption of two flavor solutions over long time scales (several hours), and consumption may offer additional insight into the multisensory operation underlying preference judgements. Finally, flavor preference judgements differ from other multisensory judgements in that they are highly subjective, and it remains unknown to what degree the integrative operations underlying preference judgements are generalizable across flavor qualities. In particular, flavors with positive and negative hedonic value are processed by partly distinct neural circuits (X. Chen, Gabitto, Peng, Ryba, & Zuker, 2011; Peng et al., 2015; Wang et al., 2018) and therefore may follow different integrative principles.

We hypothesized that multisensory flavor preference judgements adhere to the MLE framework. Alternatively, given their unique nature, the multisensory principles underlying hedonic flavor judgments may only partially adhere to the MLE framework or differ between acceptance and rejection decisions. To test this hypothesis, we evaluated the validity of the MLE framework for explaining flavor preference and consumption of both palatable and unpalatable flavor mixtures in rats. We demonstrate that multisensory preferences for palatable flavors are in line with MLE in that they can be described as a weighted average of the component judgement, and that this operation is independent of amounts consumed. We further show that—contrary to the prediction of the MLE framework—the reliability of multisensory flavor preferences does not increase relative to the component judgments. Finally, we show that hedonic judgments of unpalatable flavor mixtures follow a distinct integrative principle where components enhance each other in informing rejection decisions.

## Methods

### Animals

A total of 44 Long-Evans rats (n=24 females) were used for this study. Pregnant dams were purchased from Charles River Laboratories (www.criver.com) and litters were kept together until weaning on postnatal day 21 (P21). Rats were pair-housed after weaning and separated into single housing conditions at the start of experimental procedures, around P65. Animals were always kept on a 12-hour light-dark cycle (lights on at 6 a.m.) with *ab libitum* access to rat chow and plain water unless otherwise specified. All procedures were conducted in accordance with the NIH Guide for the Care and Use of Laboratory Animals and approved by the Institutional Animal Care and Use Committee of Wake Forest University School of Medicine.

### Stimuli

Stimuli consisted of aqueous solutions of taste and/or odor compounds (obtained from www.fishersci.com and www.sigmaaldrich.com; >98% purity, dissolved in purified water). Unisensory taste stimuli consisted of two concentrations of sucrose (10 mM and 40 mM) and citric acid (5 mM and 10 mM). Amyl acetate (0.025% weight/volume), a banana-like odor, served as the unisensory odor stimulus. At this concentration, amyl acetate has no known gustatory or trigeminal chemesthetic qualities (Rebello, Kandukuru, & Verhagen, 2015; Slotnick, Westbrook, & Darling, 1997). Multisensory stimuli consisted of binary combinations of unisensory stimuli. Plain water served as a control stimulus.

### Behavioral testing

Preference testing consisted of a series of daily two-bottle free choice consumption tests. During each daily test, animals had access to two bottles, placed side-by-side: one containing plain water; the other containing a flavor stimulus. One group of animals (n=22; n=12 females) was exposed to 5 flavor conditions featuring sucrose at two concentrations as the taste condition (see Table 1). A separate group of animals (n=12; n=5 females) was exposed to 5 flavor conditions featuring citric acid at two concentrations as the taste condition. A third group of animals (n=10; n=7 females) was exposed to 3 flavor conditions featuring citric acid at a single concentration as the taste condition. Each condition was presented 8 times in blocks, with each block featuring each condition (i.e., 5 or 3 days/block; 40 or 24 days total). The order of conditions was randomized within each block. Water and flavor bottles alternated position daily to control for side biases. Daily testing sessions occurred from 4 p.m. to 8 a.m. and lasted 16 hours. Each bottle contained 150 ml of solution ensuring animals had *ab lib* access to solutions throughout the testing session. Between testing sessions, animals had *ab lib* access to water to ensure they were not in any way restricted in their access to fluids for the entire duration of the experiment.

**Table 1.** Overview of stimulus conditions with sucrose as the taste component (Group 1)

|  | **Bottle A** |  | **Bottle B** |
|---|---|---|---|
| **Condition** | ***Taste component*** | ***Smell component*** |  |
| Low Taste | 10 mM Sucrose | - | Water |
| High Taste | 40 mM Sucrose | - | Water |
| Odor | - | 0.025% Amyl acetate | Water |
| Low Mixture | 10 mM Sucrose | 0.025% Amyl acetate | Water |
| High Mixture | 40 mM Sucrose | 0.025% Amyl acetate | Water |

**Table 2.** Overview of stimulus conditions with citric acid as the taste component (Group 2)

|  | **Bottle A** |  | **Bottle B** |
|---|---|---|---|
| **Condition** | ***Taste component*** | ***Smell component*** |  |
| High Taste | 10 mM Citric acid | - | Water |
| Odor | - | 0.025% Amyl acetate | Water |
| High Mixture | 10 mM Citric acid | 0.025% Amyl acetate | Water |

**Table 3.** Overview of stimulus conditions with citric acid as the taste component (Group 3)

|  | **Bottle A** |  | **Bottle B** |
|---|---|---|---|
| **Condition** | ***Taste component*** | ***Smell component*** |  |
| Low Taste | 5 mM Citric acid | - | Water |
| High Taste | 10 mM Citric acid | - | Water |
| Odor | - | 0.025% Amyl acetate | Water |
| Low Mixture | 5 mM Citric acid | 0.025% Amyl acetate | Water |
| High Mixture | 10 mM Citric acid | 0.025% Amyl acetate | Water |

### Data analysis

Consumption was measured by weighing bottles before and after each testing session. Preference for flavor solutions (P_F_) was calculated as flavor consumption (C_F_) relative to water consumption (C_W_) as:

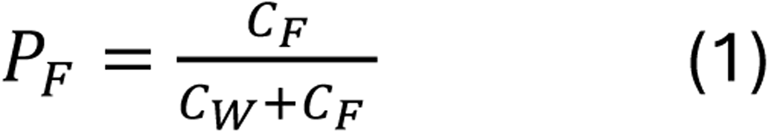

where a score >0.5 indicates a preference for the flavor solution, and a score <0.5 indicates a preference for water. Preference analyses were based on the mean and variance across repetitions of the same condition for each animal. Our primary hypothesis is that mixture preferences can be explained by Maximum Likelihood Estimation (MLE) (Ernst & Banks, 2002). According to the MLE framework for cue combination, multisensory judgements (in this case: mixture preferences [P_Mix_]) are a weighted average of the unisensory component judgements (taste preference [P_T_] and odor preference [P_O_]):

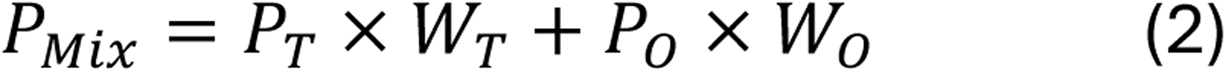

Where the weights (W) are proportional to the relative reliability of the unisensory components (A and B), as quantified by their relative variance (s^2^):

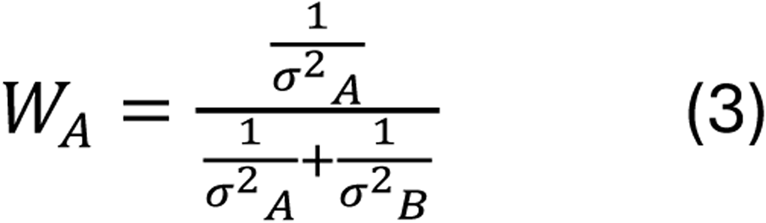

A second prediction of the MLE framework is that judgments of multisensory stimuli are more reliable (i.e., less variable) than judgments of their unisensory components:

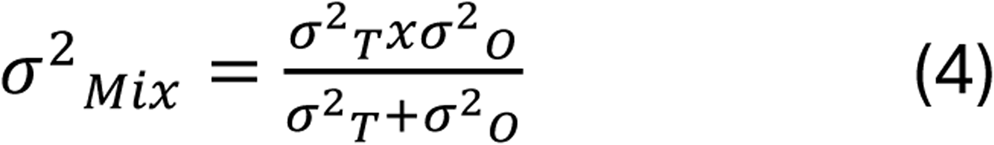

Because preferences are bound between 0 and 1, they bear a non-linear relation with respect to consumption at high preferences. To eliminate this non-linearity as a confounding factor, we predicted mixture preferences by averaging at the level of the amounts consumed, rather than the preferences themselves:

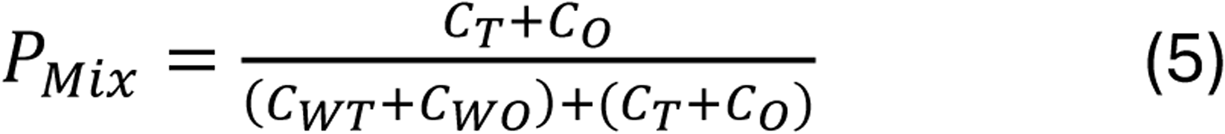

Where C indicates the amount consumed from flavor (C_T_, C_O_) and corresponding water (C_WT_, C_WO_) bottles.

Multisensory enhancement (E) was quantified as:

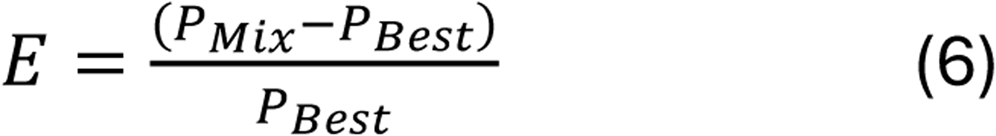

Where P_Best_ is the most effective (i.e., least preferred) unisensory component (Miller, Pluta, Stein, & Rowland, 2015; Stanford, Quessy, & Stein, 2005).

In a total of n=26 testing sessions (1.6% of total), preferences could not reliably be determined due to spillage or animals drinking the entire volume of liquid in one of the bottles, and preferences were not included in the analysis.

### Statistics

All quantities were compared across conditions using parametric tests as described in the results section (t-test, ANOVA; all two-tailed; α=0.05). Repeated measures were used whenever appropriate. Relations between quantities were evaluated using linear regression or correlation. To test the specificity of the variance-based prediction for mixture preferences, we performed a bootstrap procedure where we correlated mixture preferences with predictions obtained from variances that were randomized random pairs of unisensory component variances (one taste, one odor). This procedure was repeated 10000 times to obtain a 95% confidence interval against which the correlation coefficient obtained from the observed data was compared.

### Overlap with previously published data

A subset of animals from Group 1 of the present paper (n=12; 6 females) formed a behavioral dataset that was published previously (Allar et al., 2025).

## Results

We aimed to understand how taste and odor components of flavor are integrated to inform multisensory preference decisions. To this end, a first group of animals (n=22) underwent a series of two-bottle tests in which me measured preference (relative to plain water) for taste+smell mixture solutions as well as taste-only and odor-only component solutions featuring sucrose (a palatable sweet taste) as the taste stimulus and amyl acetate (a novel fruity smell) as the odor component (Group 1, see Table 1). We experimentally manipulated the reliability of the taste component by varying sucrose concentration (10 and 40 mM), analogous with extant manipulations of visual stimulus reliability by varying stimulus strength. To ensure a manageable number of experimental conditions, we used only a single odor stimulus and relied on individual animal variation to obtain a range of relative component reliabilities. In line with previous studies, we quantified component reliability as the inverse of the variance in component preference judgments: that is, how consistently is a flavor component used to inform a preference judgment? To estimate mean preference and variance in preference, we repeatedly measured preferences in all conditions for each animal (n=8 repetitions of n=5 conditions, for a total of n=40 conditions). Mean and variance of unisensory component preference for each animal were then used to evaluate the MLE framework for explaining mixture preference. Note that we will use “preference” to refer to the quantity obtained from comparing amounts consumed from the two bottles (eq. 1). Flavors will be referred to as “appetitive” and “aversive” based on whether the preference value for these flavor was above or below 0.5, respectively.

### Mixture preference depends on component reliability

Figure 1A shows mean preferences across animals in the 5 flavor conditions (3 unisensory and 2 multisensory). In the unisensory conditions, sucrose was generally appetitive (i.e., preferred over water), and preference increased with increasing sucrose concentration, as expected from a palatable taste (t-test comparing preference for low and high taste only to 0.5: t_21_=12.29, p<0.001 and t_21_=84.21, p<0.001, respectively; t-test comparing high and low concentration taste only: t_21_=7.59, p<0.001). Amyl acetate was generally aversive (i.e., water was preferred over amyl acetate; t-test comparing preference for odor only to 0.5: t_21_=3.34, p<0.01). Given that the odor stimulus had no *a priori* meaning, animals could in theory ignore it and simply optimize sucrose consumption. However, at both sucrose concentrations, mixture preferences were intermediate to their component preferences demonstrating that animals use both taste and smell components when expressing preferences for taste+smell mixtures (t-test comparing preference for mixture to taste only: t_21_=6.44, p<0.001 [low] and t_21_=3.31, p<0.01 [high]; t-test comparing preference for mixture to odor only: t_21_=8.83, p<0.001 [low] and t_21_=15.60, p<0.001 [high]). With respect to the exact integrative operation, our primary hypothesis is that mixture preferences can be described by the MLE framework. According to MLE, multisensory judgments are a weighted average of the component judgements, where the weights are determined by the reliability of the components. The finding presented above that mixture preferences are intermediate to component preferences is consistent with this framework. In line with previous work on other multisensory systems, reliability was quantified as the inverse of the variance of the unisensory judgments (i.e., the consistency with which unisensory component stimuli inform a perceptual judgment). To test the validity of the MLE framework, we exploited the fact that unisensory solutions ranged in reliability. First, variances in preference exhibited natural individual variation across the population of animals. In addition, we experimentally varied reliability of the taste component by presenting sucrose at two different concentrations (low: 10 mM and high: 40 mM). Figure 1B shows the variance in preference across all animals for all conditions. On average, variances in the low concentration taste only conditions were higher than in the high concentration taste only condition (t-test: t_21_=4.96, p<0.001); variances in the odor only condition were consistently lower than in the high taste only condition (t-test: t_21_=5.19, p<0.001) and statistically indistinguishable from variances in the low taste only condition (t-test: t_21_=0.31, p=0.76). To test whether component variance can explain mixture preferences on an individual mixture basis, we predicted mixture preferences using the MLE model (eq. 2) where mixture preferences are an average of the component preferences, weighted by their relative reliability (i.e., the inverse of their variance; eq. 3). Figure 1C shows mixture preferences predicted by MLE as a function of observed mixture preferences (note that this analysis combines high and low taste concentration conditions, n=44). MLE predictions were tightly correlated with observed preferences (r=0.88, p<0.001), and were statistically indistinguishable (t_43_=1.14, p=0.26). To directly evaluate the role of variance in explaining component weight, we performed a bootstrap procedure in which variances within condition were shuffled between animals. That is, predictions were made using the mean unisensory preferences from each animal, randomly paired with variances obtained from different animals in matching conditions (see Methods for detail). Predictions of mixture preference based on matched variances were significantly better than predictions of mixture preference based on random variances (Figure 1D; p<0.01).

**Figure 1.**
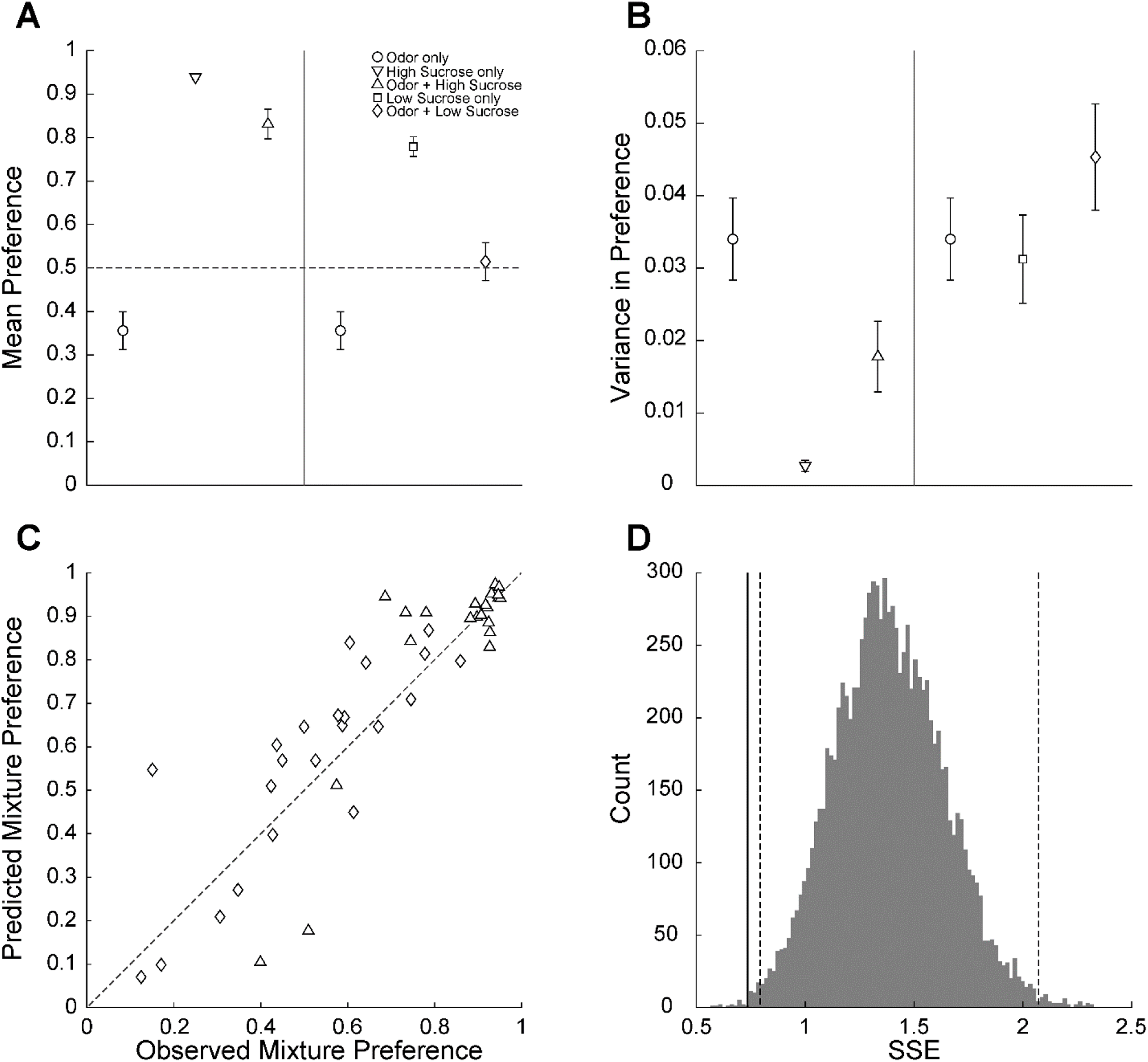
(A) Mean±SEM preference relative to water for all five conditions across all animals tested with sucrose (Group 1; n=22). Vertical line separates high concentration (left) and low concentration (right). Dashed horizontal line indicates indifference (preference=0.5). (B) Mean±SEM variance in preference for all five flavor conditions (n=22). (C) MLE-predicted mixture preference as a function of observed mixture preference for all animals in both low and high sucrose concentration conditions (n=22). Dashed line indicates x=y. (D) Distribution of SSE values from the bootstrap procedure in which component variances were randomly shuffled between animals (n=10000 iterations). Dashed vertical lines indicate the 95% confidence intervals, solid vertical line indicates SSE obtained from observed (matched) variances.

### Relation between preference and consumption

The findings presented above suggest that component variance plays a critical role in informing mixture preferences. However, the use of preference as a readout of hedonic value presents a potential confounding factor in that it is calculated from flavor consumption as a proportion of the total amount consumed and is bound between 0 and 1. As a result, preference values saturate as consumption increases. Figure 2A shows the relation between flavor preference and flavor consumption, revealing a strong non-linearity at high consumption values. As a result, any change in consumption relative to the unisensory taste condition caused by mixing in the odor would have a smaller effect on preference at high compared to low concentration taste components. That is, an equal reduction in consumption in high and low concentration mixture conditions (relative to taste only) would by definition produce preferences that appear more strongly weighted toward more preferred taste components. Such an effect could thus potentially explain our findings without taking into account variance or reliability. To directly test this, we determined how taste only consumption is affected by the addition of an odor component. Figure 2B shows amounts consumed from both bottles in all conditions. Addition of an odor component to the flavor bottle reduced consumption relative to the taste only condition, and increased consumption from the corresponding water bottle. However, changes in consumption from the mixture conditions relative to the corresponding taste only conditions were not equal for high and low concentrations (t-test comparing the change in consumption from taste only to mixture conditions between low and high concentrations: flavor bottle: t_21_=2.32 p<0.05; water bottle: t_21_=4.18. p<0.001), indicating that adding an odor did not change consumption in an additive manner. We also predicted mixture preferences based on a linear sum of component consumption (eq. 5). Figure 2C shows mixture preferences predicted by arithmetic averaging of consumption values. The model consistently underestimates high concentration mixture preferences (t-test: t_21_=4.87, p<0.001) and overestimates low concentration mixture preferences (t-test: t_21_=2.70, p<0.05). Moreover, the degree to which this model under/overestimated preferences did not correlate with the amount of flavor consumption (low concentration mixtures: r=0.03, p=0.90; high concentration mixtures: r=0.15, p=0.50). Thus, a simple additive model at the level of flavor consumption cannot explain the observed pattern in preferences.

**Figure 2.**
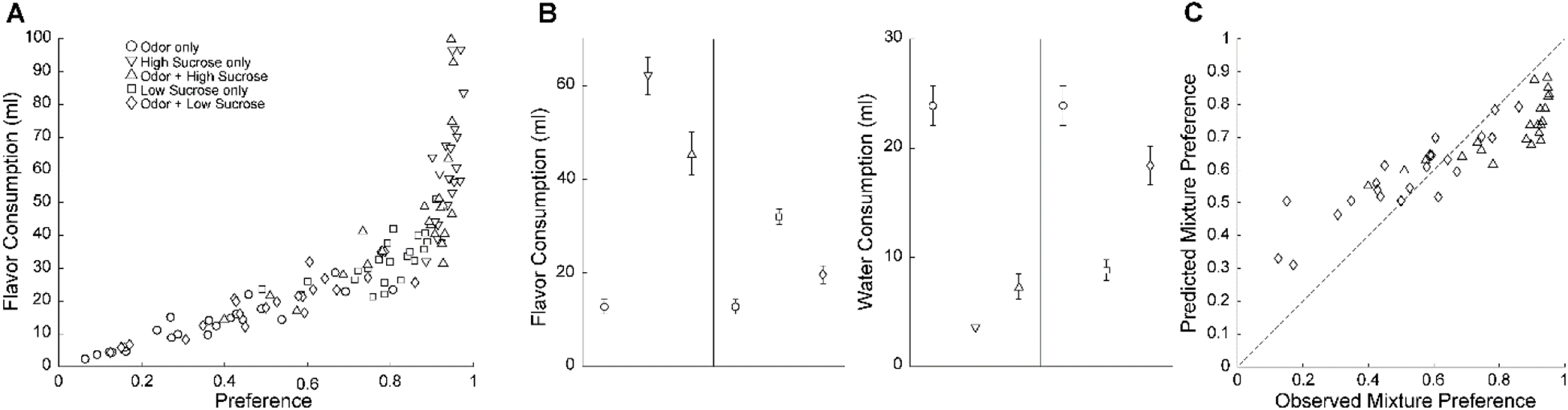
(A) Flavor bottle consumption (ml) as a function of flavor preference for all animals across all five conditions in Group 1 (n=22). Each datapoint represents a single animal in a single condition. (B) Mean±SEM flavor consumption (left) and corresponding water consumption (right) for all conditions (n=22). Vertical lines separate high (left) and low (right) concentrations. (C) Mixture preference predicted by arithmetic averaging of component consumption values (eq. 5) as a function of observed mixture preference for all animals in both mixture conditions (n=44). Dashed line indicates x=y.

### The effect of multisensory integration on the reliability of multisensory judgments

The results presented above demonstrate that mixture preference judgements follow the first prediction of the MLE framework: multisensory judgements are a weighted average of the component judgements, and the weights of the components depend on their relative reliability. The second prediction of the MLE framework states that reliability-dependent cue weighting increases the reliability of multisensory judgements relative to their component judgements. Contrary to this prediction, Figure 1B shows that the variance in preference in mixture conditions is increased relatively to the taste component conditions. Indeed, Figure 3 shows that the MLE model consistently underestimates the variance of mixture preferences (eq. 4; t-test comparing predicted and observed variances: t_43_=6.03, p<0.001).

**Figure 3.**
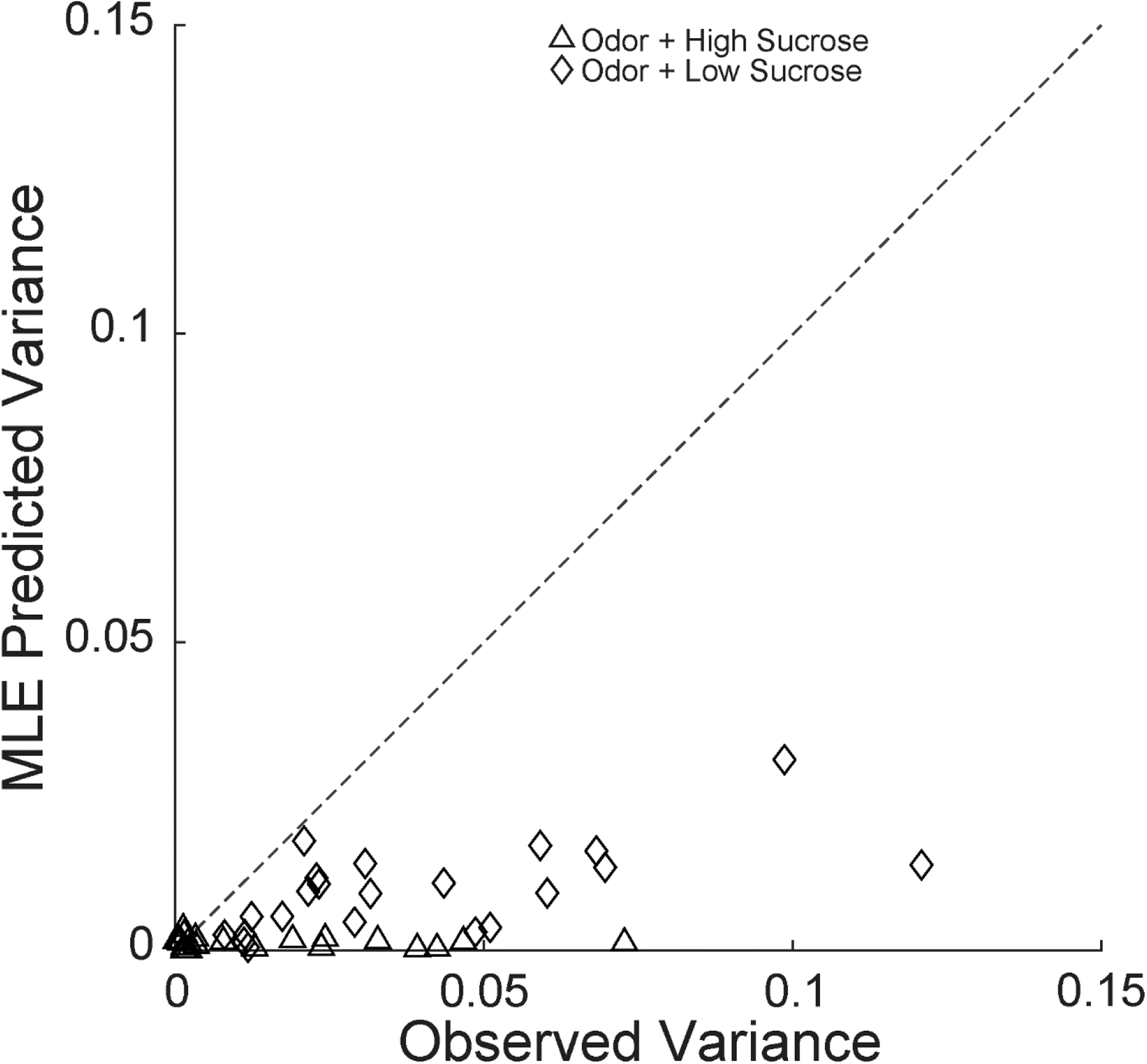
MLE-predicted variance in mixture preference (eq. 4) as a function of observed variance in mixture preference for all animals in both low and high concentration conditions in Group 1 (n=44). Dashed line indicates x=y.

### Stability of multisensory integration over the course of testing

Our design involves repeated testing on the same conditions over the course of multiple weeks. It is therefore possible that preferences for individual stimulus component or the way components are integrated undergoes experience-dependent changes. Notably, previous work has shown that odors readily become associated with tastes when presented in mixture. Figure 4A-E shows taste and odor component preferences as a function of testing block. Linear regression of preference on block revealed that preferences were stable over the course of the experiment for all conditions (R^2^ ranging from 0.01 to 0.50, all p>0.29). To assess the stability of multisensory integration, we compared preferences for mixtures and their components for each testing block. Figure 4F shows mixture-component differences for mixtures featuring low and high concentrations of sucrose. Linear regression of difference in preference on testing block revealed that multisensory interactions were stable over the course of the experiment for all components (R^2^ ranging from 0.02 to 0.17, all p>0.10). Thus, preferences for unisensory components, as well interactions between components, were stable across the duration of the experiment.

**Figure 4.**
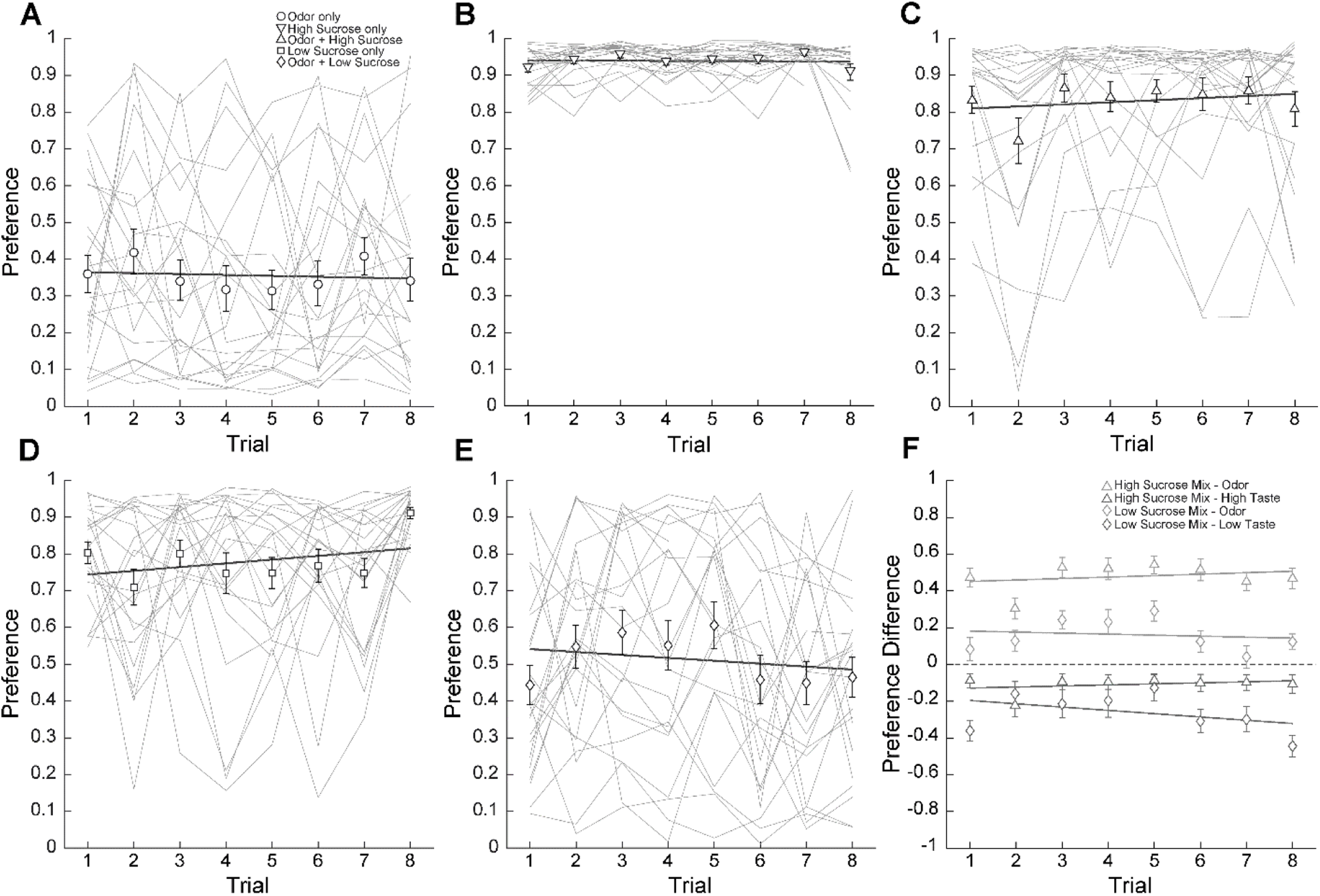
(A-E) Mean±SEM preference relative to water (black symbols) for each condition as a function of testing block for all animals in Group 1 (n=22). Gray lines indicate individual animals. Solid black line indicates linear regression fit. Conditions shown: (A) High sucrose only, (B) Odor only, (C) Odor+high Sucrose, (D) Low sucrose only, (E) Odor+low Sucrose. (F) Mean ± SEM difference in preference between each mixture condition and the corresponding taste (dark gray symbols) and odor (light gray symbols) unisensory component conditions as a function of testing block. Solid lines indicate linear regression fits. Dashed horizontal line indicates no difference.

### Integration of same-value flavor cues

The results presented above provide evidence for the idea that reliability-dependent cue combination underlies hedonic judgements of flavor mixtures. However, these results were obtained from unisensory components that have opposite valence: sucrose was generally appetitive; amyl acetate generally aversive. Large differences between component preferences allowed us to reliably test the MLE model for cue combination. However, sucrose and amyl acetate can be considered incongruent with respect to hedonic value, and the data do therefore not demonstrate generalizability to mixtures of components with the same hedonic value. To address this, we first evaluated the validity of the MLE model for a subset of animals that preferred amyl acetate relative to water (n=4). Figure 5A shows preferences for one of these animals in all conditions, revealing a pattern that is generally consistent with the one observed across the population. Note that under these conditions, the range of possible intermediate preference values is greatly reduced. Figure 5B shows observed mixture preferences and mixture preferences predicted by MLE separately for all 4 animals in both high and low concentration mixture conditions (n=8 conditions total). Preferences were accurately predicted by the MLE model (r=0.90, p<0.01; t-test comparing observed and predicted preferences: t_7_=0.05, p=0.96), demonstrating that reliability-dependent weighting holds for mixtures of taste and odor components that both have positive valence.

**Figure 5.**
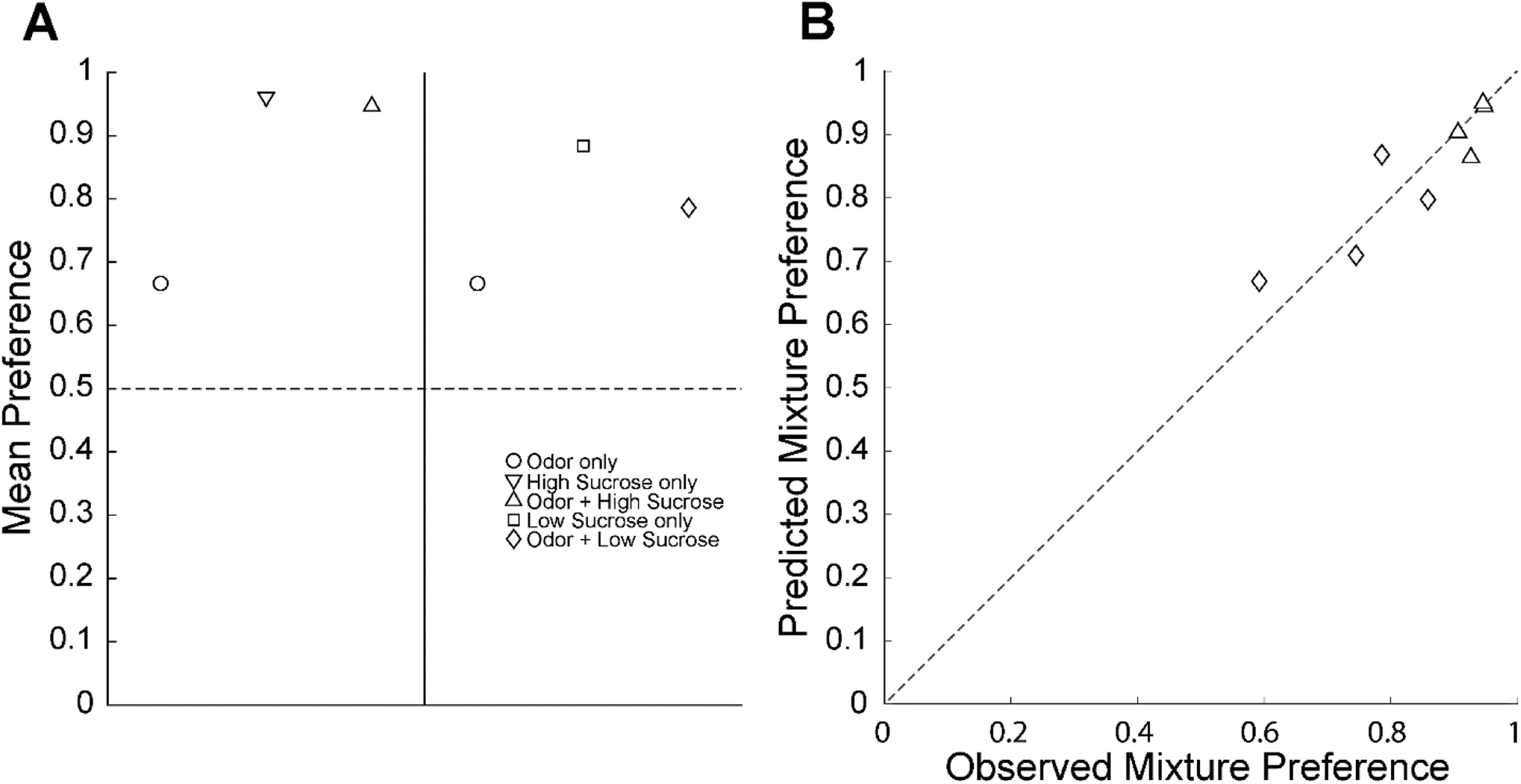
(A) Mean preference relative to water for all five conditions for a representative animal that preferred amyl acetate over water. Vertical line separates high concentration (left) and low concentration (right) conditions. Dashed horizontal line indicates indifference (preference=0.5). (B) MLE-predicted mixture preference as a function of observed mixture preference for all animals that preferred amyl acetate over water (n = 4) in high and low sucrose concentration mixture conditions (n=8). Dashed line indicates x=y.

In a separate set of animals (n=22), we tested preferences for mixtures containing amyl acetate and an unpalatable taste (citric acid). One subset of these animals (Group 2, n=10) were tested on 3 conditions, featuring 10 mM citric acid as the taste stimulus. The other animals (Group 3, n=12) were tested on 5 conditions, featuring a low (5 mM) and a high (10 mM) concentration of citric acid as taste stimuli. The results from these two groups will be presented together in the analyses presented below (i.e., data on 5 mM citric acid will include 12 animals; data on 10 mM citric acid will include 22 animals). Figure 6A shows mean preferences across animals in the 5 conditions. As expected, both citric acid and amyl acetate were generally aversive, and aversion increased with increasing citric acid concentration (t-test comparing taste only to 0.5: 5mM [low], 10mM[high]; t-test comparing odor only to 0.5: t_11_=5.82, p<0.001 and t_21_=15.9, p<0.001; t-test comparing low and high concentration taste only conditions: t_32_=1.96, p=0.06). In contrast to the pattern of results obtained from the experiments using sucrose as a taste stimulus, preferences for taste-smell mixtures featuring citric acid as a taste stimulus were not intermediate to the unisensory component preferences. Thus, judgements of citric acid-amyl acetate are not consistent with the MLE framework. Figure 6B shows observed mixture preferences as a function of mixture preferences predicted by MLE. MLE significantly underestimates mixture preferences (t-test comparing observed versus predicted mixture preferences: t_33_=3.96, p<0.001). Instead, the results appear consistent with the enhancement framework for multisensory integration. This framework states that components of a multisensory stimulus interact to enhance intensity or detectability (B. E. Stein & Stanford, 2008). In the present context, this framework predicts that aversions to mixture solutions are stronger than aversions to the most aversive component solution. We tested this prediction by first determining—for each animal and each mixture—what the most aversive component was (i.e., taste or smell). In high concentration conditions, taste was the most aversive component in n=17/22 cases (77.3%); in low concentration conditions, taste was the most aversive component in n=6/12 cases (50%). Figure 6C shows mixture preferences as a function of the most aversive component. Mixtures were significantly more aversive than the most aversive component (t_33_=2.60, p<0.05), suggesting that the components of citric acid-amyl acetate mixtures interact to enhance aversiveness. We also tested a second prediction of the enhancement framework, namely that the amount of enhancement is greatest when the unisensory components are least effective in isolation (the principle of inverse effectiveness). In the present context, this principle predicts that the increase in aversiveness for mixtures relative to their components is strongest for less aversive components. Figure 6D shows the degree of enhancement (eq. 6) as a function of preference for the least preferred component. The amount of enhancement increases with decreasing unisensory aversiveness. Finally, we assessed the stability of component preferences and multisensory interactions as we did above for mixtures containing sucrose. Figure 7A-E shows preferences for the unisensory component solutions across testing block; Figure 7F shows differences in preference between mixtures and their most aversive component across testing blocks. Linear regression did not reveal systematic changes in preference for unisensory solutions across blocks (R^2^ ranging from 0.00 to 0.03, all p>0.12). However, we did find a significant decrease in preference for the odor only condition after the first testing block (t-test comparing odor only preference between blocks 1 and 2: t_21_=2.59, p<0.05). Figure 7F shows that despite this change in preference for the odor component from block 1 to block 2, the pattern of multisensory interactions was stable across the experiment (R^2^ ranging from 0.00 to 0.03, all p>0.12). Taken together, these findings demonstrate that taste-smell integration underlying hedonic judgements of citric acid-amyl acetate mixtures differs qualitatively from hedonic judgements of sucrose-amyl acetate mixtures.

**Figure 6.**
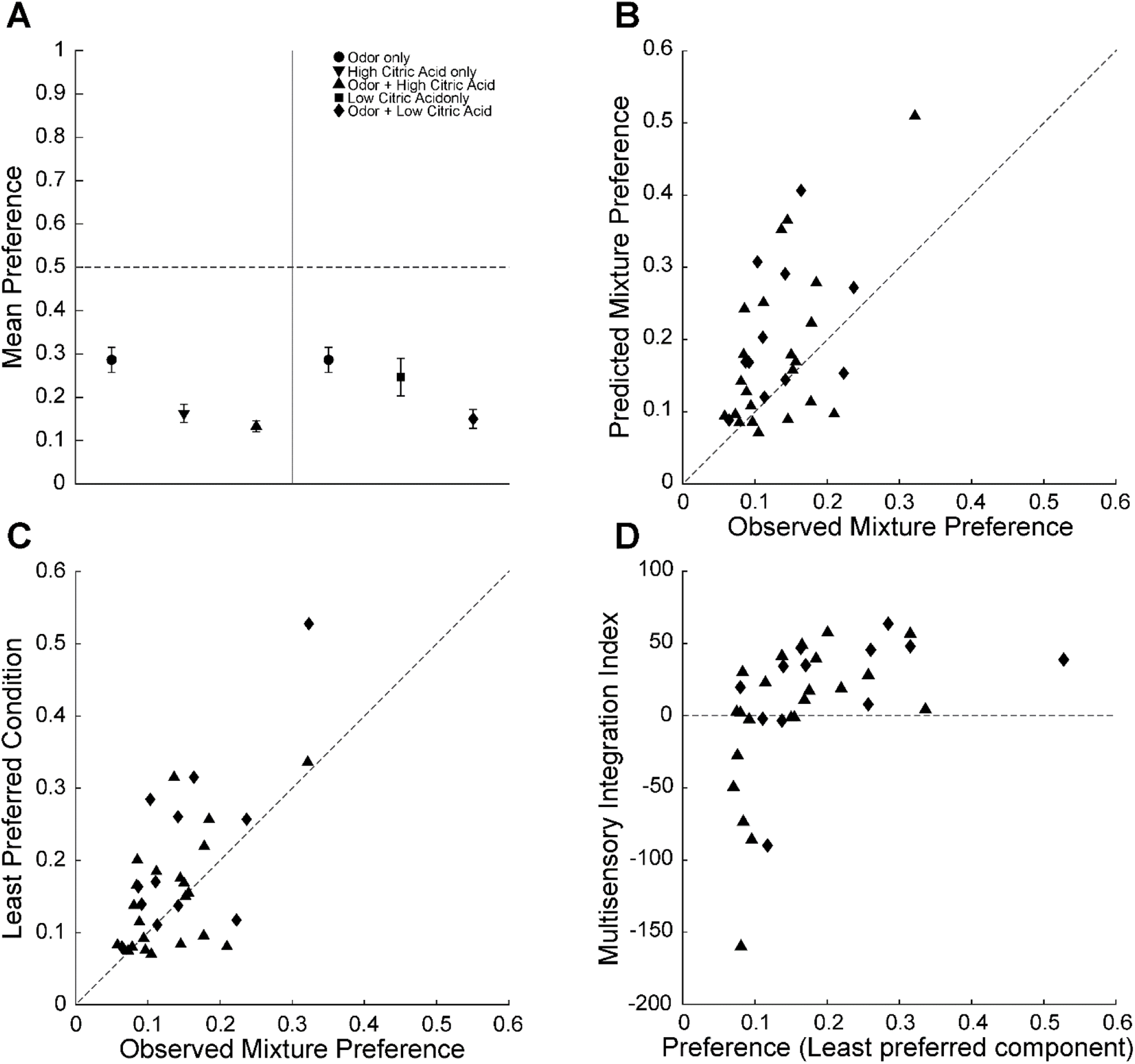
(A) Mean±SEM preference relative to water for all five conditions across all animals tested with citric acid as the taste stimulus (n=22 for high concentration conditions [Group 2, 3]; n=12 for low concentration conditions [Group 2]). Vertical line separates high concentration (left) and low concentration (right) conditions. Dashed horizontal line indicates indifference (preference=0.5). (B) MLE-predicted mixture preference as a function of observed mixture preference for all animals in both low and high citric acid concentration mixture conditions (n=34). Dashed line indicates x=y. (C) Least preferred unisensory component preference as a function of observed mixture preference for all animals in both mixture conditions (n=34). Dashed line indicates x=y. (D) Multisensory enhancement index (eq. 6) as a function of preference for the least preferred unisensory component condition (n=34). Dashed horizontal line indicates no enhancement.

**Figure 7.**
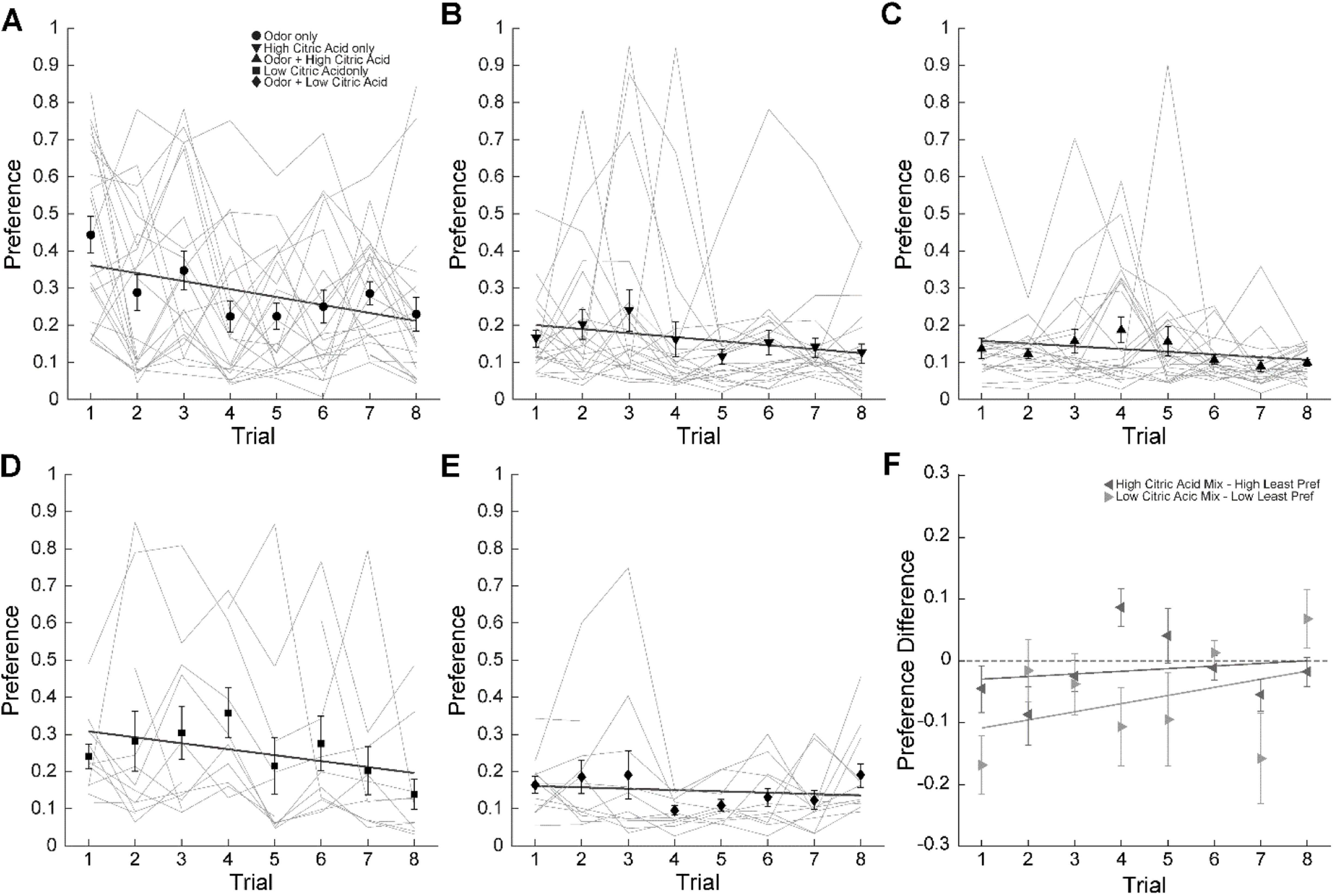
(A-E) Mean±SEM preference relative to water (black symbols) for each condition as a function of testing block for animals tested with citric acid as the taste stimulus (n=22 for high concentration conditions; n=12 for low concentration conditions). Gray lines indicate individual animals. Solid black line indicates linear regression fit. Conditions shown: (A) High citric acid only, (B) Odor only, (C) Odor+high citric acid, (D) Low citric acid only, (E) Odor+low citric acid. (F) Mean±SEM difference in preference between each mixture condition and its least preferred unisensory component as a function of testing block. Solid lines indicate linear regression fits. Dashed horizontal line indicates no difference.

## Discussion

Our results show that multisensory flavor preferences (relative to water) for mixtures containing a palatable taste (sucrose) reflect a weighted average of the component preferences, where the weights of the components depend on their relative reliability. That is, the reliability with which taste and odor components inform preference decisions determines how much weight they carry in informing preferences for a mixture of taste and odor components. This finding is consistent with a previous report from our lab (Allar et al., 2025) that used a partly overlapping dataset (see Methods) and further extends the insight into taste-smell integration toward hedonic flavor judgments in several ways. Two findings suggest that mixture preferences cannot be explained by an additive operation, where the hedonic values of the components are simply added to yield mixture preferences. First, a subset of animals that preferred both taste and odor components over water did not express mixture preferences that were higher than the unisensory component preferences. Instead, mixture preferences for these animals were consistent with reliability-dependent weighting. However, amyl acetate was appetitive only in a small subset of our animals. Future work using generally appetitive odor stimuli across the population will more effectively test whether reliability-dependent weighting holds for mixtures of preferred odor and taste components. Second, mixture consumption did not reflect a sum of taste and smell component consumption, and an additive model did not accurately predict observed mixture preferences. Instead, component reliability played a critical role in determining multisensory flavor preferences regardless of component preference magnitude.

Reliability-dependent integration is a highly adaptive operation in that less reliable sensory signals are ignored more effectively when making judgements in a multisensory context. A similar operation has been described in other multisensory domains. For example, humans weight haptic and visual cues in a reliability-dependent manner when estimating the size of an object (Ernst & Banks, 2002), and (non-)human primates combine vestibular and visual cues in a similar manner when judging the direction of self-motion (Butler et al., 2010; Fetsch et al., 2009). In these studies, multisensory judgements have been shown to be consistent with predictions made by the maximum likelihood estimation framework. The finding discussed above that mixture preferences reflect a reliability-dependent weighted average of the component preferences is consistent with this framework. However, our findings are not consistent with a second key prediction of the MLE framework: that mixture judgements are themselves more reliable than component judgements. Indeed, the observed reliability of mixture preferences was consistently lower than predicted by MLE. One possible explanation lies in the fact that flavor preference judgments are not made on a trial-by-trial basis as in typical psychophysical studies on multisensory integration. Instead, preferences are derived from a dynamic sampling process that unfolds over long timescales (∼14 hours in the present study). Previous efforts to characterize the microstructure of flavor sampling revealed that different dynamics contribute to consumption, and eventually preference. For example, preferences obtained from two-bottle testing are driven not only by hedonic aspects of the solutions (i.e., liking), but also by motivation (i.e., wanting) as well as learning and memory (Ballintyn, Ksander, Katz, & Miller, 2023; Berridge, Robinson, & Aldridge, 2009; Verharen, de Jong, Zhu, & Lammel, 2023). Moreover, these dynamics appear to be driven by at least party distinct neural systems that each explain variation in preference between individual animals and different contexts (Castro & Berridge, 2014; Verharen et al., 2023; Wulff, Cooper, Kodjo, Abel, & Thompson, 2023). Detailed analysis of sampling behavior in taste, odor and mixture conditions could provide insight into how these different processes contribute to variation in preference. For example, mixture conditions may differ from unisensory conditions in appetitive aspects of behavior (e.g., how often does an animal visit the flavor bottle versus the water bottle), or consummatory aspects of behavior (e.g., how much does an animal consume from a bottle during each visit to that bottle). Another possibility is that consummatory responses in mixture conditions do not reflect a weighted average of the component conditions, but that animals display modality switching (Ernst & Bülthoff, 2004; Meijer et al., 2019; Wozny, Beierholm, & Shams, 2010), where the amount consumed during each visit equals either odor or taste consumption. Finally, it is possible that appetitive and/or consummatory aspects of behavior change over the course of testing in different ways during mixture and unisensory conditions.

The second main finding from the present study is aversion of mixtures containing unpalatable taste and odor components are enhanced relative to aversions to the components in isolation. This enhancement was strongest when the components were least informative. Such a multisensory operation has been described in the literature on multisensory stimulus detection. For example, concurrent presentation of visual and auditory stimuli increases perceived intensity relative to the components in isolation (Lippert, Logothetis, & Kayser, 2007) and improves stimulus localization (B. E. Stein, Huneycutt, & Meredith, 1988; B. E. Stein, Meredith, Huneycutt, & McDade, 1989) and identification (Werner & Noppeney, 2010). Such enhancement effects are strongest when the unisensory components are weakest. Similar findings have been reported in the context of flavor, where the addition of an odor increases detectability of a subthreshold concentration of taste (Dalton, Doolittle, Nagata, & Breslin, 2000; Shepard, Veldhuizen, & Marks, 2015; Veldhuizen, Shepard, Wang, & Marks, 2010). It is possible that enhanced rejection of unpalatable mixtures in the present study is the result of enhanced detectability/intensity. The observed integrative pattern could be adaptive in that it lowers the intake of potentially harmful substances. A remaining question is whether the observed enhancement is due to the unpalatable nature of the taste component specifically. Future studies using unpalatable taste and palatable odor components will test this idea.

Recent work has started to elucidate the neural substrates underlying taste-smell integration. Notably, neurons in the insular primary gustatory cortex of rodents respond to taste and smell inputs (Allar et al., 2025; Maier, 2017; Samuelsen & Fontanini, 2017; Stocke & Samuelsen, 2024), and human imaging studies observed overlapping activation patterns evoked by the two modalities in the insular cortex (de Araujo, Rolls, Kringelbach, McGlone, & Phillips, 2003; Khorisantono, Veldhuizen, & Seubert, 2025; Veldhuizen, Nachtigal, Teulings, Gitelman, & Small, 2010). A recent study from our lab demonstrated that spiking activity of neurons in the GC of rats reflects reliability-dependent weighting of multisensory inputs (Allar et al., 2025). The neural pathways underlying multisensory enhancement of flavor aversion remain unknown. One possibility is that aversive flavor inputs are processed by unique neuronal populations within GC that perform different integrative operations. This would be consistent with the idea that palatable and unpalatable tastes are represented in spatially distinct subregions of GC (X. Chen et al., 2011; Peng et al., 2015; Wang et al., 2018). However, given a wealth of evidence that palatable and unpalatable tastes are represented in overlapping neuronal populations within GC (K. Chen, Kogan, & Fontanini, 2021; Katz, Simon, & Nicolelis, 2001; Levitan et al., 2019; Vincis, Chen, Czarnecki, Chen, & Fontanini, 2020), it is also possible that enhancement of multisensory flavor inputs occurs outside of GC in circuits that specifically mediate flavor avoidance behaviors, like the amygdala.

Our results were obtained from animals that were naïve to the flavor stimuli at the start of the experiment. The observed stability in integrative patterns over the course of testing further suggests that the ability to integrate taste and smell components of flavor does not depend on experience, consistent with pervious work in rats (Elliott & Maier, 2020; Hua et al., 2024). However, the extant literature suggests that multisensory flavor perception is experience-dependent in several ways. First, experiencing an odor in mixture with a palatable/unpalatable taste can result in an increase/decrease in preference for that odor when presented in isolation. Such conditioning effects has been observed in both rodent (Christensen, Triplett, & Maier, 2022; Holman, 1975; McQueen, Fredericksen, & Samuelsen, 2020; Sclafani & Ackroff, 1994) and human (R. J. Stevenson, R. A. Boakes, & J. P. Wilson, 2000; R. J. Stevenson, Boakes, & Prescott, 1998; Richard J Stevenson, Robert A Boakes, & Judith P Wilson, 2000; R. Stevenson & Boakes, 2004; R. Stevenson, Prescott, & Boakes, 1995; Yeomans, Mobini, Elliman, Walker, & Stevenson, 2006) studies. In the present experiment, animals repeatedly experienced amyl acetate in mixture with sucrose or citric acid over the course of preference testing. The fact that we did not observe appetitive conditioning in the sucrose group could be the result of latent inhibition and/or memory extinction. Because preference testing sessions were relatively long in duration, consumption of amyl acetate in isolation during unisensory conditions may have prevented amyl acetate from becoming associated with sucrose during mixture conditions, in line with latent inhibition. Any association between amyl acetate and sucrose may also extinguish rapidly over the course of consumption during odor only conditions. A similar lack of conditioning effects has been observed previously under similar conditions (i.e., overnight flavor versus water two-bottle preference testing) (Colbert et al., 2022). Second, several studies have shown that taste-smell interactions depend on flavor congruency (Amsellem & Ohla, 2016; Fondberg et al., 2018; Schifferstein & Verlegh, 1996; Seo et al., 2013). For example, enhancement of flavor detection and discrimination only occurs when taste and smell components had been experienced together in mixture (Cai, Allar, & Maier, 2024; Shepard et al., 2015). As discussed above, the integrative patterns observed in the present study occur in the absence of any detectable congruency between taste and odor components in the mixtures, and previous studies in rats have failed to observe effects of congruency on multisensory flavor preferences. However, it is possible that integrative patterns change after extensive experience with specific taste-odor combinations. Future studies that experimentally manipulate flavor experience before the onset of testing or exploit possible innate taste-odor congruencies (Tieman et al., 2012) will further provide further insight into this question.

## Acknowledgements

Many thanks to Chad Collins for help with data collection, and to Paul Miller for helpful suggestions on analysis of consumption data. This work was supported by NIH NIDCD R01 DC020212 (JXM).

